# AURKA controls oocyte spindle assembly checkpoint and chromosome alignment by HEC1 phosphorylation

**DOI:** 10.1101/2024.11.01.621527

**Authors:** Cecilia S. Blengini, Robert J. Mendola, G. John Garrisi, Jason E. Swain, Karen Schindler

## Abstract

In human oocytes, meiosis I is error-prone, causing early miscarriages and developmental disorders. The Aurora protein kinases are key regulators of chromosome segregation in mitosis and meiosis and their dysfunction is associated with aneuploidy. Oocytes contain three Aurora kinase (AURK) proteins, but only AURKA is necessary and sufficient to support oocyte meiosis in mice. However, the unique molecular contributions of AURKA remain unclear. Here, using a combination of genetic and pharmacological approaches, we evaluated how AURKA phosphorylation regulates outer kinetochore function during oocyte meiosis. We found that the outer kinetochore protein Ndc80/HEC1 is constitutively phosphorylated at multiple residues by Aurora kinases during meiosis I, but that Serine 69 is specifically phosphorylated by AURKA in mouse and human oocytes. We further show that Serine 69 phosphorylation regulates spindle assembly checkpoint (SAC) activation and chromosome alignment during meiosis I. These results provide a fundamental mechanistic understanding of how AURKA regulates meiosis and kinetochore function to ensure meiosis I fidelity.

## Introduction

Creation of haploid gametes, eggs and sperm, through meiosis is essential for species propagation. It is therefore critical that meiosis occurs accurately, yet meiosis I is highly error prone in human oocytes. These errors lead to early miscarriages and developmental disorders (Gruhn et al., 2019; Hassold et al., 2007). Understanding how meiosis I is regulated is therefore critical for determining why human eggs are uniquely prone to chromosome segregation errors.

The Aurora protein kinases are key regulators of chromosome segregation in mitosis and meiosis (Carmena and Earnshaw, 2003; Nguyen and Schindler, 2017). Unlike mitotically dividing cells which express two Aurora kinases (AURKA and AURKB), oocytes express three (AURKA, AURKB and AURKC). Because it is meiosis-specific, much work has focused on determining AURKC functions in oocytes (Balboula et al., 2016; Balboula and Schindler, 2014; Cairo et al., 2023; Fellmeth et al., 2015; Quartuccio et al., 2017; Schindler et al., 2012; Vallot et al., 2018; Yang et al., 2010; Yang et al., 2015). However, our works show that AURKA is the only AURK that is necessary and sufficient for mouse oocyte meiosis (Blengini et al., 2021a; Blengini and Schindler, 2024; Blengini et al., 2024; Nguyen et al., 2018; Schindler et al., 2012). In determining the spatio-temporal functions of AURKA, we observed that neither targeting AURKA to the acentriolar microtubule organizing centers (aMTOCs) at spindle poles nor targeting it to the chromosomes fully rescued the spindle and chromosome alignment defects in oocytes lacking all three *Aurk*s (Blengini et al., 2024). We speculated that an additional population of AURKA, either at kinetochores or on spindle microtubules, contribute to spindle elongation and chromosome alignment.

Kinetochore-microtubule (KT-MT) attachments are critical for chromosome alignment and spindle structure. The kinetochore is a complex protein structure that is the interface between centromeres and spindle MTs, regulating MT attachment, MT dynamics and chromosome movement. The NDC80 complex is part of the outer kinetochore, and it is the direct link between KTs and MTs (Cheeseman et al., 2006; DeLuca and Musacchio, 2012). The NDC80 complex is composed of four proteins: SPC24 and SPC25, which interface with the inner kinetochore and HEC1 and NUF2 which bind MTs (Musacchio and Desai, 2017). HEC1 undergoes precise phospho-regulation to ensure KT-MT dynamics allowing cycles of attachment and detachment until KT-MT stabilization is achieved (DeLuca et al., 2006). The HEC1 N-terminus contains nine phosphorylation sites (Ser4, Ser5, Ser8, Ser15, Ser44, Thr49, Ser55, Ser62 and Ser69) (DeLuca et al., 2006; DeLuca et al., 2011; Santaguida and Musacchio, 2009). These sites are important for regulating KT-MT affinity thereby controlling correction of erroneously attached MTs, and they are critical for controlling chromosome oscillation and movement during mitosis (DeLuca et al., 2011; Zaytsev et al., 2015; Zaytsev et al., 2014). In mitosis, AURKA and AURKB phosphorylate HEC1, but the specific contribution of each AURK differs. For example, inhibition of AURKA and AURKB in RPE-1 and HeLa cells showed that Ser55 is phosphorylated by both kinases (DeLuca et al., 2011; DeLuca et al., 2018; Iemura et al., 2021), whereas Ser44 is primarily phosphorylated by AURKB (DeLuca et al., 2011; DeLuca et al., 2018), and Ser69 is primarily phosphorylated by AURKA (DeLuca et al., 2018). Based on the pattern of phosphorylation and the phenotype when each phosphorylation site was altered, different functions for each site were assigned. Although the function of pSer44 is not well understood, pSer55 is critical for destabilization of KT-MT attachments during early prometaphase (DeLuca et al., 2011; DeLuca et al., 2018) and for chromosome oscillation at metaphase (Iemura et al., 2021). pSer69 is also important for chromosome oscillation at metaphase decreasing the probability of lagging chromosomes (DeLuca et al., 2018; Iemura et al., 2021). In mouse oocytes, HEC1 phosphorylation is important for chromosome alignment (Yoshida et al., 2015), for determining the length of the meiosis I spindle and for restricting aMTOCs at spindle poles (Courtois et al., 2021; Gui and Homer, 2013). However, it is not known how HEC1 phosphorylation is regulated, and which Aurora kinase is involved in fine-tuning this regulation during mouse oocyte meiosis.

To understand the regulatory landscape of HEC1 during oocyte meiosis, we used a combination of genetic and pharmacological approaches to determine the pattern of HEC1 phosphorylation. Furthermore, because oocytes express AURKA, AURKB and AURKC, we determined the role of each AURK in phosphorylating HEC1 residues. Compared to mitosis, we find differences in meiosis where full phosphorylation of Ser44 requires other unknown kinases, whereas phosphorylation of Ser55 and Ser69 are AURKC and AURKA specific, respectively. Given the importance of AURKA in oocyte biology, we determine the role of Ser69 phosphorylation and discovered that it is not only conserved in human oocytes, but that is important for regulating spindle assembly checkpoint (SAC) activation and chromosome alignment in mouse oocytes. These results add a new layer of mechanistic understanding of the essential roles of AURKA in regulating kinetochore phosphorylation in oocyte meiosis.

## Results

### Serines 44, 55 and 69 of HEC1 are constitutively phosphorylated during oocyte meiotic maturation

In mitosis, the N-terminus of HEC1 is highly phosphorylated during pro-metaphase and this phosphorylation decreases as the cells approach metaphase. However, it is not known how HEC1 phosphorylation behaves during meiotic maturation in mammalian oocytes. Of the nine HEC1 phosphorylation sites, antibodies that specifically detect five sites (serines 8, 15, 44, 55 and 69) exist. Therefore, we first evaluated which sites are phosphorylated in mouse oocytes. We detected phosphorylation at three serine residues: 44, 55 and 69 (Fig 1A); phosphorylation of serines 8 and 15 was not detectable above background signal. Next, to determine the temporal pattern of HEC1 phosphorylation during meiotic maturation, we obtained WT mouse oocytes at early pro-metaphase I, late-pro-metaphase I and metaphase I and detected these three phosphorylated residues. Although we found a modest, but statistically significant reduction of pSer55 at metaphase I, the three sites remained constitutively phosphorylated throughout these phases of meiotic maturation and dephosphorylation did not occur at metaphase I (Fig 1B,C). These data suggest that HEC1 phosphorylation and regulation is different in mouse oocyte meiosis I compared to mitosis where they are dephosphorylated as the cell cycle progresses. There are differences between mitosis and meiosis I that could explain these different temporal patterns. In mitosis, the tension between sister chromatids promotes spatial separation of AURKB from its KT substrates. This separation is coordinated with the recruitment of phosphatases to KTs to stabilize the KT-MT attachments via HEC1 dephosphorylation (Foley et al., 2011; Kabeche and Compton, 2013; Lampson and Cheeseman, 2011). Because of the unique meiosis I bivalent chromosome geometry, this separation does not occur even when bivalents are stretched. Therefore, HEC1 and other AURKC KT-substrates remain phosphorylated until localized PP2A-B56 phosphatase activity rises enough to counteract AURKC activity (Yoshida et al., 2015). Therefore, the intra-KT stretching phase and stabilization of KT-MT attachments are temporally separated (Davydenko et al., 2013; Kitajima, 2018; Yoshida et al., 2015). It is possible that constitutive HEC1 phosphorylation is advantageous, allowing oocytes more time to correct abnormal MT attachments. Alternatively, KT-MT activities may occur with an intermediate level of attachment stability to allow attachment but high turnover. These models require further evaluation.

**Figure 1.**
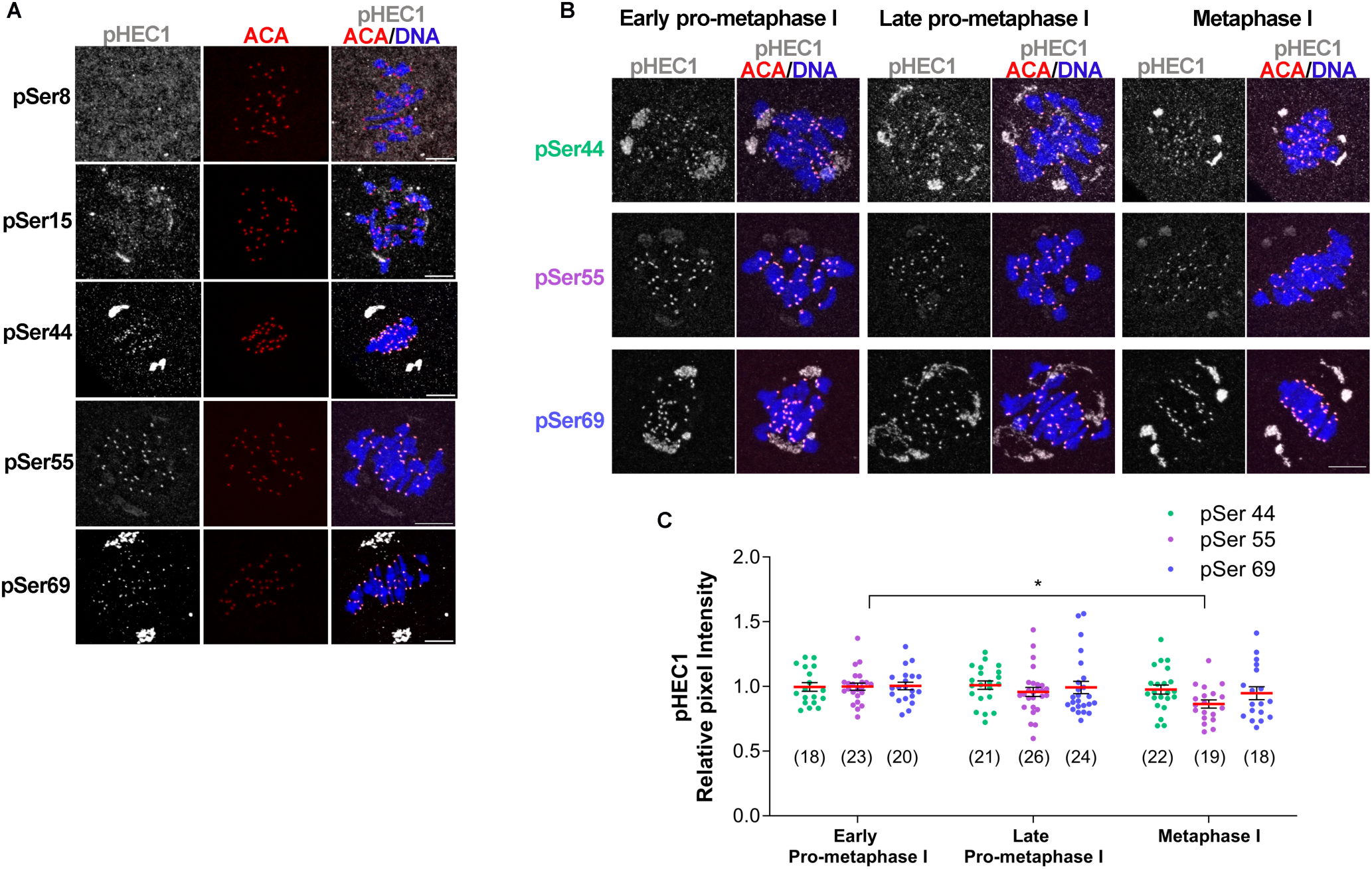
Temporal pattern of HEC1 N-terminus phosphorylation. (A) Representative confocal images of WT oocytes stained with phospho-specific antibodies to detect phosphorylated Serines (pSer) 8, 15, 44, 55 and r69 (gray), ACA to label kinetochores (red) and DAPI to detect DNA (blue). (B) Representative confocal images of WT oocytes matured to early -prometaphase I, late pro-metaphase I and metaphase I that were stained with phospho-specific antibodies to detect pSer44, pSer55 and pSer69 (gray), ACA to label kinetochores (red) and DAPI to detect DNA (blue). Scale bar: 10µm. (C) Quantification of phosphorylated HEC1 at pSer44 (green dots), pSer55 (purple dots) and pSer69 (blue dots) in (B). The numbers of oocytes analyzed are in brackets from 2 independent experiments. One-way ANOVA pSer44: p= 0.7698; pSer55 * p<0.05; pSer69: p=0.6585.

### Aurora kinases A and C phosphorylate HEC1 at Serine 55 and 69 in mouse oocytes

Next, we determined the requirement of the Aurora kinases in HEC1 phosphorylation in mouse oocytes. To address this question, we evaluated the levels of HEC1 phosphorylation at Serines 44, 55 and 69 by immunocytochemistry in metaphase I oocytes lacking all three AURKs (ABC-KO). Confirmation of this knockout mouse strain was reported previously (Blengini et al., 2024). We observed that pSer55 and pSer69 were nearly absent in ABC-KO oocytes compared to WT oocytes (Fig 2C-F), suggesting that at least one AURK is critical for the phosphorylation of these residues. However, pSer44 was only reduced by ∼50% in ABC-KO oocytes compared to WT oocytes (Fig 2A, B). These data indicate that another kinase also phosphorylates Ser44.

**Figure 2.**
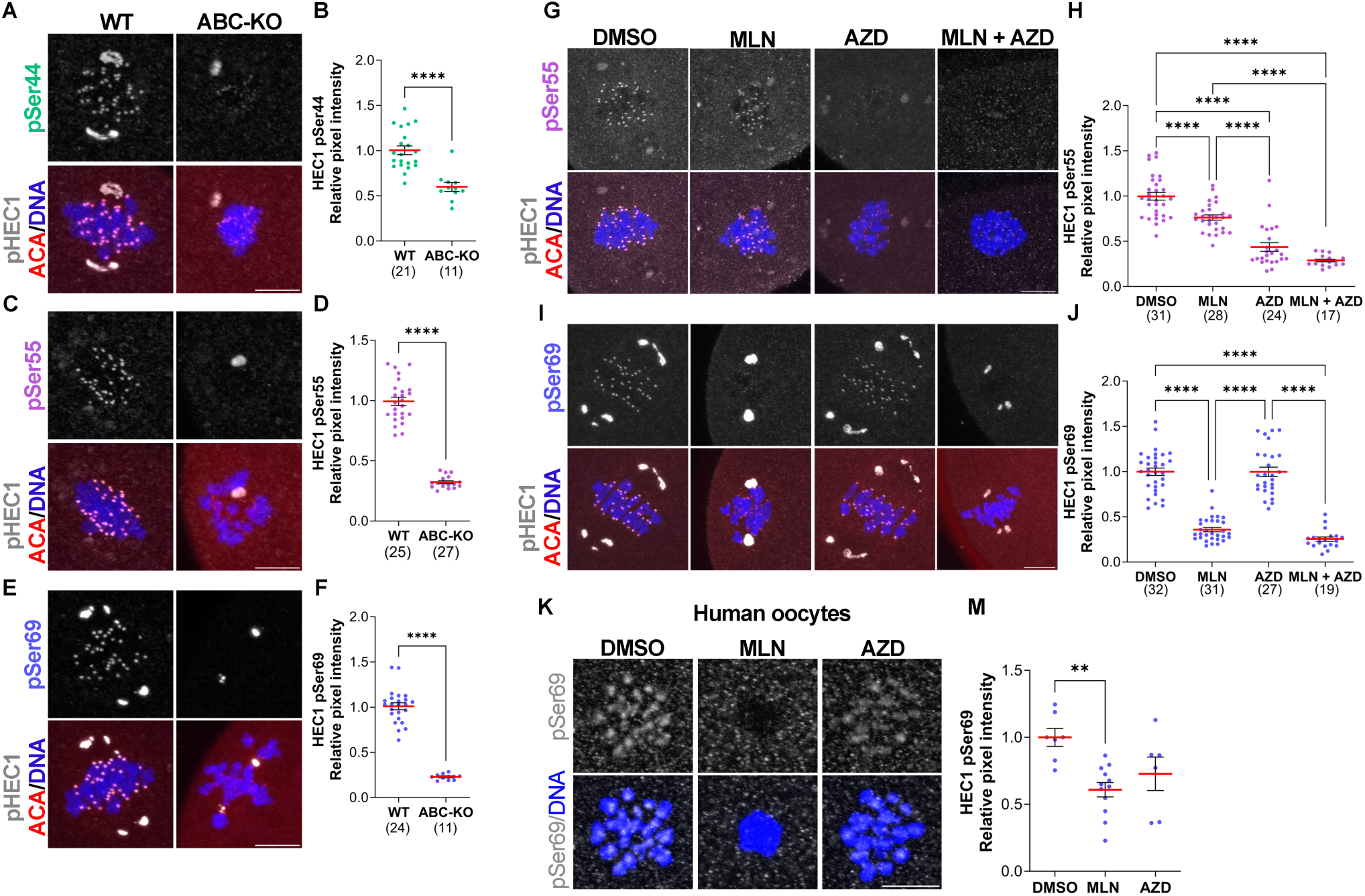
Specificity of Aurora kinase phosphorylation of HEC1 N-terminal residues. (A), (C), (E) Representative confocal images of WT and ABC-KO oocytes at metaphase I stained with phospho-specific antibodies to detect pSer44 (A), pSer55 (C) and pSer69 (E) (gray), ACA (red) and DNA (blue). (B), (D), (F). Quantification of phosphorylated HEC1 at pSer44 (B) (green dots) in (A), pSer55 (D) (purple dots) in (C) and pSer69 (F) (blue dots) in (E). Students T-test, **** p<0.0001. (G), (I) Representative confocal images of WT oocytes matured to metaphase I and treated with DMSO (control), MLN, AZD or MLN + AZD and stained with phospho-specific antibodies to detect pSer55 (G) and pSer69 (I) (gray), ACA (red) and DNA (blue). (H), (J) Quantification of phosphorylated HEC1 at pSer55 (H) (purple dots) in (G) and pSer69 (J) (blue dots) in (I). One-way ANOVA: **** p<0.0001. (K) Representative confocal images of human oocytes matured to metaphase I and treated with DMSO (control), MLN or AZD stained with a phospho-specific antibody to detect pSer69 (gray). DNA was detected with DAPI (blue). (M) Quantification of pSer69 (blue dots) in (K). One-way ANOVA, ** p<0.01. Scale bar: 10µm. The number of oocytes analyzed is in brackets.

Next, to evaluate AURK specificity in Ser55 and Ser69 phosphorylation, we used a pharmacological approach applying MLN8237 (MLN) and AZD1152 (AZD) small molecule inhibitors to WT oocytes at concentrations with demonstrated specificity for AURKA or AURKB/C, respectively (Blengini et al., 2022). These experiments were performed at metaphase I to acutely inhibit the kinases. Upon AURKA inhibition, pSer55 signal reduced by 25%, whereas upon AURKB/C inhibition pSer55 signal reduced by 60% (Fig. 2G, H). Inhibition of AURKA/B/C did not significantly reduce pSer55 signal more than AURKB/C inhibition alone. Taken together, the data suggests that Serine 55 is primarily phosphorylated by AURKC in WT mouse oocytes. pSer69 was absent upon inhibition of AURKA and no additional reduction was observed when all three kinases were inhibited (Fig. 2I, J) suggesting that, as in somatic cells, Ser69 is primarily phosphorylated by AURKA.

Because the AURKs can compensate for one another (Aboelenain and Schindler, 2021; Blengini et al., 2021b; Blengini and Schindler, 2024; Blengini et al., 2024; Nguyen et al., 2018), we took a genetic approach to determine the compensatory ability of AURKs to phosphorylate these HEC1 sites. We generated single and double AURK knockout mouse oocytes and evaluated the levels of Ser55 and Ser69 phosphorylation at metaphase I. pSer55 was significantly reduced by 25% in oocytes lacking *Aurkc* (C-KO and AC-KO)(Fig. S1A, B; Table S1). The difference of pSer55 immunoreactivity between C-KO and AC-KO was not significantly different. These results suggest that in absence of AURKC, AURKA/B cannot phosphorylate Ser55. pSer69 was strongly reduced in all mouse strains lacking AURKA in oocytes (Fig. S1C, D; Table S1) and Serine 69 phosphorylation was most significantly reduced in AC-KO oocytes (Fig. S1C, D; Table S1). These genetic data suggest that AURKC can partially compensate in A-KO oocytes and phosphorylate Ser69.

These results revealed some similarities and differences in how the N-terminus of HEC1 is regulated between mitosis and mammalian oocyte meiosis. For example, Ser44 is less AURKB/C specific in meiosis compared to mitosis and less AURKA/B compensation occurs at S55 in meiosis compared to mitosis. However, Ser69 is predominantly phosphorylated by AURKA in both mitosis and oocyte meiosis. Interestingly, AURKB does not appear to phosphorylate any of these sites in oocytes (Fig. 2, Table S1), likely because of the expression of AURKC, the *Aurk* homolog with highest homology to *Aurkb*, predominates at kinetochores. The expression of the third AURK member promotes a division of tasks during oocyte meiotic maturation, induced by changes in localization and expression levels of each AURK isoform that could affect the pattern of HEC1 phosphorylation. However, AURKA in oocytes localizes similar to somatic cells because AURKA is localized predominantly to spindle poles and has a minor chromosomal population (Kratka et al., 2022). Therefore, AURKA-dependent phosphorylation of Ser69 in oocytes, is conserved between cell types.

Given the critical functions of AURKA in mouse oocytes, we further investigated whether HEC1 pSer69, which is solely dependent on AURKA, is phosphorylated in human oocytes. To answer this question, we obtained discarded, cryopreserved prophase I-arrested human oocytes and matured them *in vitro* with the MLN and AZD AURK inhibitors. First, Ser69 phosphorylation was detectable in metaphase I human oocytes and this mark was localized at kinetochores (Fig. 2K). Second, we asked if AURKA is responsible for Ser69 phosphorylation in human oocytes as it is in mouse. We found that pSer69 was significantly reduced upon AURKA inhibition (MLN) and reduced, but not significantly, upon AURKB/C inhibition (AZD) (Fig. 2K, M). These results suggest that phosphorylation of HEC1-Ser69 by AURKA is conserved between mouse and human oocytes, but that there may be some redundancy with AURKB/C in human that we do not observe in mouse.

### Phosphorylation of HEC1-Ser69 is important for spindle assembly checkpoint activation and chromosome alignment at metaphase I

Given the importance of AURKA in oocyte biology and the conservation of HEC1-Ser69 phosphorylation between mouse and human oocytes, we decided to further to investigate the role of pSer69 during mouse oocyte meiotic maturation. We created HEC1 constructs that encoded either a non-phosphorylated HEC1 variant where Ser69 was mutated to an alanine (69A) or a phospho-memetic HEC1 variant where Ser69 was mutated to an aspartic acid (69D). RNA from these constructs were injected into WT mouse oocytes which were then cultured for the time it takes control oocytes to reach metaphase II. We found that most oocytes expressing either WT HEC1 or the 69D variant arrested in metaphase I and failed to reach metaphase II. In contrast, 20% of the oocytes expressing the 69A variant progressed past metaphase I, and reached metaphase II (Fig. 3A, B). As a control we observed that the expression levels of GFP fusion proteins were comparable between the different constructs (Fig. 3C, D), indicating that the phenotypic differences are not because of different expression levels. These results suggest that Ser69 phosphorylation contributes to cell-cycle regulation in oocytes.

**Figure 3.**
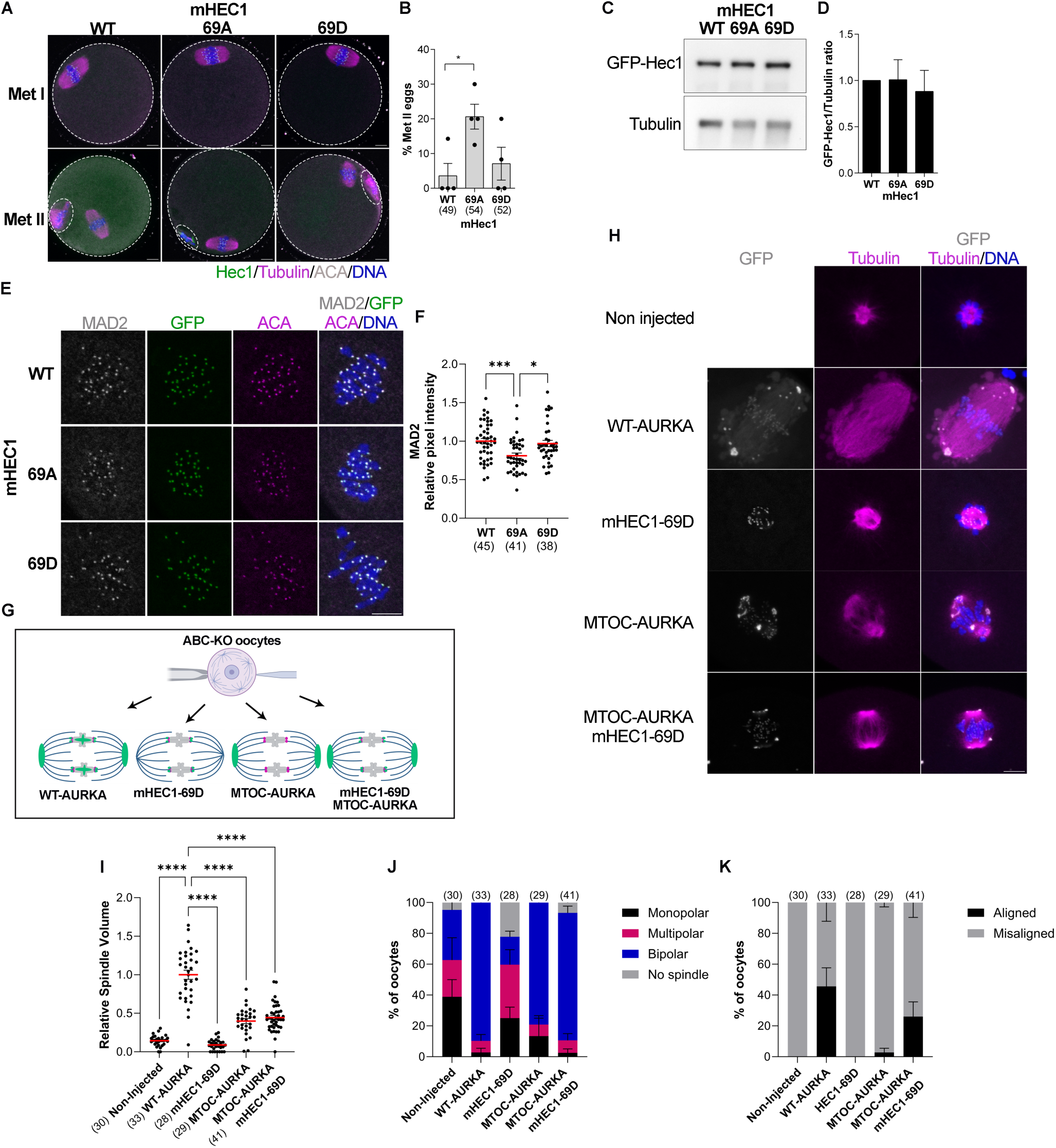
HEC1-Ser69 phosphorylation regulates SAC activation and chromosome alignment. A) Representative confocal images of WT oocytes expressing the indicated *Hec1* mRNAs, matured to metaphase II stained with Tubulin to detect the spindle (pink), and DAPI to detect DNA (blue). Localization of HEC1-GFPs is green. (B) Quantification of the percentage of oocytes at Metaphase II in (A). (C) Representative Western blot images of WT Metaphase I oocytes expressing the indicated *Hec1-Gfp* mRNAs. (D) Quantification of GFP/Tubulin ratio from (C). One-way ANOVA, p= 0.8665. (E) Representative confocal images of WT oocytes expressing the indicated *Hec1-Gfp* mRNAs at Metaphase I and stained with MAD2 (gray), ACA to detect kinetochores (pink) and DNA (blue). Localization of HEC1-GFPs is green. (F) Quantification of relative MAD2 pixel intensity from (E). One-way ANOVA, ***p< 0.001; *p<0.05. (G) Schematic of the experimental design. ABC-KO oocytes expressing different AURKA-targeted and HEC1-69D constructs shown in green. Part of this schematic was generated using BioRender. (H) Representative confocal images of Metaphase I ABC-KO oocytes expressing the indicated mRNAs. Non-injected ABC-KO oocytes were control. Oocytes were stained with tubulin to detect the spindle (green) and DAPI to detect DNA (blue). Localization of the targeted AURKA and HEC1 proteins are in gray. (I) Quantification of the relative spindle volume from (H). One-way ANOVA, **** p<0.0001. (J) Quantification of the percentage oocytes with different spindles phenotypes in (H). Two-way ANOVA. Interaction: **** p<0.0001. (K) Quantification of the percentage oocytes with chromosome misaligned in (H). Two-way ANOVA. Interaction: **** p<0.0001. Scale bar: 10µm. In brackets the number of oocytes analyzed.

Previous studies evaluating somatic cells overexpressing HEC1, showed that these cells arrested in metaphase due to spindle assembly checkpoint (SAC) activation and had high levels of MAD2 at kinetochores, a marker of SAC activation (Diaz-Rodríguez et al., 2008; Kemmler et al., 2009). To investigate the possibility that Ser69 phosphorylation is involved in regulating SAC activation, we evaluated MAD2 levels at kinetochores in metaphase I in oocytes expressing the HEC1 variants. Oocytes with reduced MAD2 recruitment have a weakened SAC and will extrude polar bodies when challenged with SAC-inducing conditions (Homer et al., 2005; Wassmann et al., 2003). Oocytes expressing HEC1-S69A had a 20% reduction in kinetochore-localized MAD2, suggesting that phosphorylation of Ser69 contributes to SAC activation in mouse oocytes (Fig. 3E, F). A caveat with our experimental approach is that the WT oocytes still express endogenous HEC1. Therefore, endogenous WT HEC1 could contribute to the observed mild phenotype in the S69A-expressing oocytes. We note that this result is consistent with previous studies where MAD2 levels were reduced in HEC1-KO oocytes (Yoshida et al., 2020). The results are also consistent with the phenotype where HEC1 KO oocytes express a mutant of HEC1 where nine N-terminal phosphorylation sites were mutated to Alanines. Those oocytes underwent premature anaphase I onset, a phenotype consistent with the SAC being prematurely satisfied or weakened (Courtois et al., 2021). Our data extends this finding to show that of the nine phosphorylation sites, phosphorylation of HEC1-Ser69 by AURKA is significantly responsible for SAC activation.

HEC1 is also important for promoting spindle bipolarization (Courtois et al., 2021; Yoshida et al., 2020) by recruiting PRC1, an antiparallel MT cross-linking protein, to kinetochores (Yoshida et al., 2020). HEC1 phosphorylation also restricts the aMTOCs to spindle poles (Courtois et al., 2021) and helps ensure chromosome alignment (Yoshida et al., 2015). In our previous study where we analyzed the localized functions of AURKA on meiotic spindle formation, we observed that neither tethering AURKA to aMTOCs nor tethering it to the chromosomes fully rescued the spindle and chromosome alignment defects in ABC-KO oocytes (Blengini et al., 2024). We speculated that we were missing a population of AURKA at kinetochores that contributes to spindle elongation and chromosome alignment. Therefore, we explored whether the AURKA-dependent phosphorylation at Ser69 could improve phenotypic rescue in ABC-KO oocytes. To this end, we overexpressed MTOC-targeted AURKA together with HEC1-S69D in ABC-KO oocytes (Fig. 3G) and examined metaphase I spindle volume, spindle bipolarity and chromosome alignment. We compared these parameters to ABC-KO oocytes expressing either WT-AURKA, MTOC-AURKA, or HEC1-S69D alone as controls. First, we evaluated whether the overexpression of these constructs could restore phosphorylation of HEC1 at Ser55 and Ser69. Compared to non-injected control ABC-KO oocytes, expression of WT-AURKA, MTOC-targeted AURKA and mHEC1-S69D either alone or in combination did not restore pSer55 in ABC-KO oocytes (Fig. S2A). This failure to restore pSer55 is consistent with a requirement for AURKC to target Ser55 in WT oocytes (Fig. 2G, H). In contrast, pSer69 was restored in ABC-KO oocytes expressing either WT-AURKA or MTOC-targeted AURKA, although to different extents (Fig. S2B-E). These results are consistent with our genetic and pharmacological results showing that Ser69 is primarily phosphorylated by AURKA in mouse oocytes (Fig. 2I, J). Moreover, the data show that AURKA localized at aMTOCs can phosphorylate Ser69 at kinetochores, although not to the same extent as WT-AURKA (Fig. S2B-C). These results are consistent with AURKA ability to phosphorylate kinetochore substrates when chromosomes are close to spindle poles in early pro-metaphase I in oocytes (Chmatal et al., 2015). Alternatively, in oocytes expressing MTOC-targeted AURKA, we observed a dim GFP signal at kinetochores (Blengini et al., 2024) which could be phosphorylating Ser69. The restoration levels of oocytes expressing MTOC-AURKA and HEC1-69D together were reduced compared to oocytes expressed WT-AURKA or MTOC-targeted AURKA alone, suggesting that expression of HEC1-S69D does not fully replace endogenous HEC1 at kinetochores in ABC-KO oocytes (Fig. S2B, E).

We next asked whether spindle parameters and chromosome alignment defects were rescued by co-expressing MTOC-targeted AURKA and HEC1-S69D. ABC-KO oocytes have small spindle volumes, 25% bipolarity, and 100% chromosome misalignment (Fig. 3H-K). Consistent with our previous findings, ABC-KO oocytes expressing WT-AURKA had restored metaphase I spindle volumes, 90% spindle bipolarity, and 50% chromosome alignment. When HEC1-S69D was expressed, the phenotype was similar to ABC-KO non-injected control oocytes: small spindle volumes, spindles with mixed polarity, and 100% chromosome misalignment (Fig. 3H-K). When MTOC-targeted AURKA was expressed, spindle volumes were partially restored, but, although 80% of the spindles were bipolar, nearly all of the oocytes still had chromosome misalignment (98%) (Fig. 3H-K). When MTOC-AURKA and HEC1-S69D were expressed together, spindle volumes and bipolarity were similar to oocytes expressing MTOC-AURKA alone. Interestingly, the percentage of oocytes with chromosome alignment (∼30%) was comparable to oocytes expressing WT-AURKA (50%) (Fig. 3H-K), suggesting that Ser69 phosphorylation regulates chromosome alignment in mouse oocytes. Taken together, these results suggest that oocytes require AURKA activity at spindle poles and at kinetochores during meiotic maturation. However, if there are only two discrete populations of AURKA or if a gradient of AURKA diffuses through the spindle via the liquid-like spindle domain needs further investigation.

In sum, we uncovered precise, fundamental mechanisms of how two of the three AURKs contribute to the phospho-regulation of the HEC1 N-terminus in mouse oocyte meiosis. By probing the requirements of Ser69 phosphorylation we uncovered a new molecular mechanism explaining how AURKA regulates cell-cycle progression by activating the SAC and controlling chromosome alignment during oocyte maturation (Fig. 4). This mechanism is likely conserved between mouse and human.

**Figure 4.**
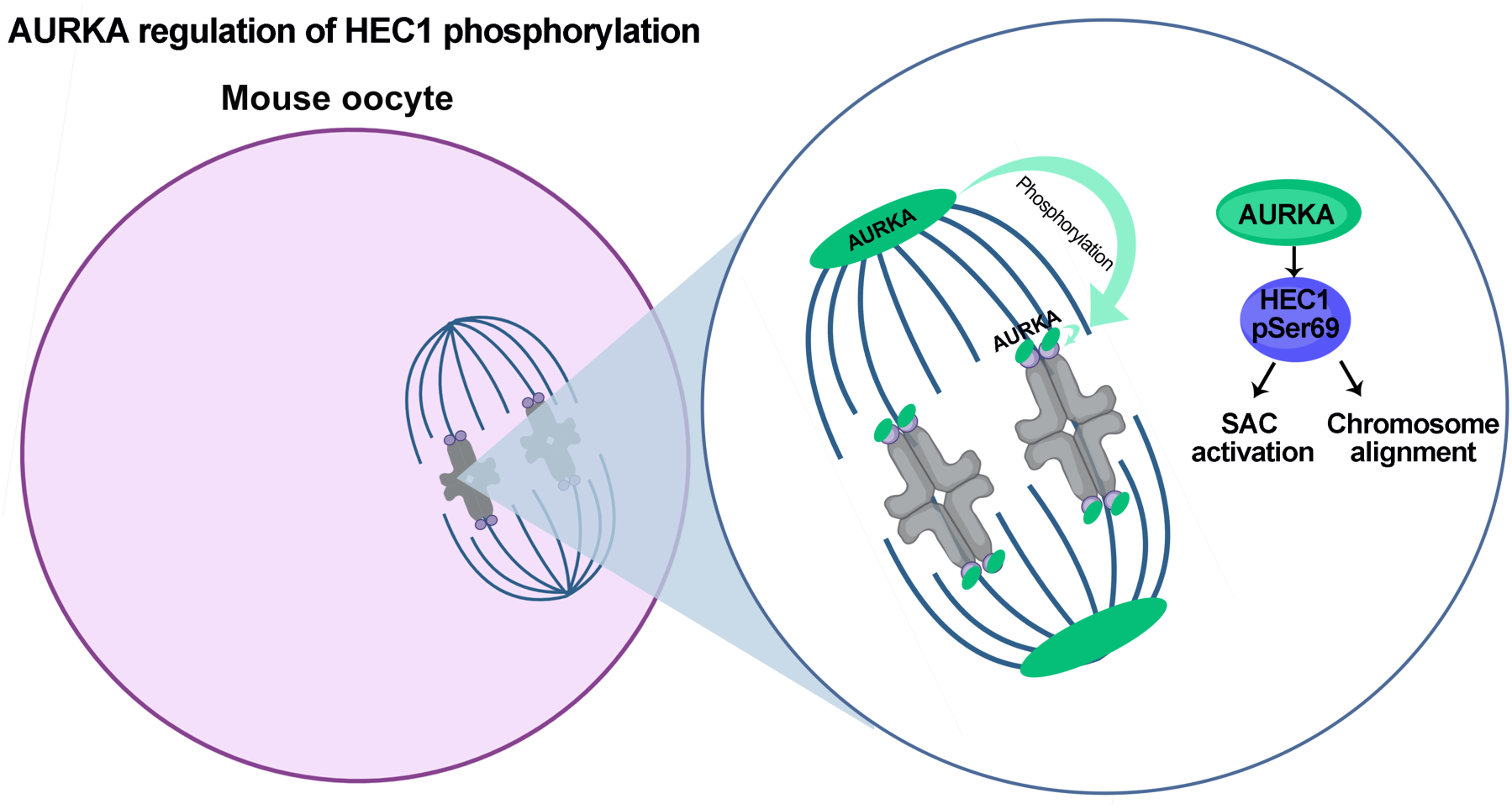
Model of specific in Aurora kinase A regulation of HEC1 N-terminus phosphorylation in oocytes. Populations of AURKA from the spindle poles and the kinetochores regulates spindle assembly checkpoint activation and chromosome alignment during oocyte maturation by the phosphorylation of kinetochore protein HEC1 at Ser69 during oocyte meiosis. Part of this figure was generated using BioRender.

## Material and Methods

### Mouse strains

To determine the temporal pattern of Hec1 phosphorylation sites and for microinjection experiments we used WT CF1 females (Envigo) aged 6-12 weeks. The knockout strains used in this work were on a C56BL/6N background: Wild type (WT) *Aurka^f^*^/f^, and *Aurkb*^f/f^ but lacking the Cre recombinase transgene; *Aurka^f^*^/f^ Gdf9-Cre (A-KO) (Blengini et al., 2021a); *Aurkb^f^*^/f^, and Gdf9-Cre (B-KO) (Nguyen et al 2018); *Aurkc*^-/-^ (C-KO) (Kimmins et al 2007; Schindler et al 2012); *Aurka^f^*^/f^, *Aurkb^f^*^/f^, and Gdf9-Cre (AB-KO) (Blengini et al., 2024), *Aurka^f^*^/f^ *Aurkc*^-/-^ and Gdf9-Cre (AC-KO); *Aurka*^*f*/f^, *Aurkb*^*f*/f^ *Aurkc^-/-^* and Gdf9-Cre (ABC-KO)(Blengini and Schindler, 2024; Blengini et al., 2024). All mice were described and validated previously. Mice were housed on a 12–12□h light-dark cycle, with constant temperature and with food and water were provided ad libitum. Animals were maintained in accordance with guidelines of the Institutional Animal Use and Care Committee of Rutgers University (protocol 201702497). All oocyte experiments were conducted using healthy female mice aged 6–16□weeks. Genotyping was performed before weaning and repeated when the animals were used for experiments as previously described (Blengini et al 2024)

### Mouse oocyte collection, maturation and microinjection

Prophase I-arrested oocytes were isolated from ovaries from females hormonally primed 48 hours earlier with 5 I.U. of pregnant mare’s serum gonadotropin (PMSG) (Lee Biosolutions #493–10). Ovaries were placed in minimal essential medium (MEM) containing 2.5 μM milrinone (Sigma-Aldrich #M4659) to prevent meiotic resumption. To induce meiotic resumption, oocytes were cultured in milrinone-free Chatot, Ziomek, and Bavister (CZB) medium in an atmosphere of 5% CO_2_□in air at 37°C. To determine the temporal pattern of HEC1 phosphorylation, oocytes were matured for different periods depending on the experimental conditions: to reach early prometaphase I, 3 hours after milrinone wash; for late prometaphase I, 5 hours after milrinone wash; for metaphase I, 7-7.5 hours after milrinone wash. For acute inhibition of Aurora kinases, oocytes were matured for 7 hours to reach metaphase I and the incubated in the respective inhibitor treatment for 3 hours. To avoid the entrance to anaphase I, we inhibited the proteasome by adding 5mM MG132 (Selleck Chemicals, #S2619) to the culture media.

For microinjection, prophase-arrested oocytes were maintained in CZB supplemented with 2.5 μM milrinone to keep them arrested. CF1 oocytes were microinjected with 200ng/μl□mouse HEC1-Gfp *(WT HEC1),* 200 ng/μl□mouse HEC1-S69A-Gfp *(HEC1-S69A),* 200 ng/μl□mouse HEC1-69D-Gfp *(HEC1 69D).* ABC-KO oocytes were injected with 100 ng/μl□*Aurka-EYfp (WT AURKA),* 100 ng/μl□CDK5FRAP fr-*Aurka-EYfp (MTOC-AURKA)* (Blengini et al., 2024), 100 ng/ μl□mouse HEC1-69D-Gfp *(HEC1 69D)or with* 100 ng/μl□CDK5FRAP fr-*Aurka-EYfp and* 100 ng/ μl□mouse HEC1-69D-Gfp together.

Microinjected oocytes were cultured overnight in CZB supplemented with 2.5 μM milrinone to allow protein expression prior to the procedures. Subsequently, the oocytes were matured for 7-7.5 hours to reach metaphase I.

### Human oocytes

Human oocytes used in this study were discarded immature oocytes (Prophase I) originating from patients undergoing routine IVF and elected oocyte cryopreservation cycles under the IRMS/CCRM-NJ IRB approved protocol (WCG Aspire protocol # #20193402). All IRB consents include patient consent to participate and publish. All human oocytes were obtained from eleven patients ranging from ages 28-37 years old, and cryopreserved using vitrification.

Cryopreserved human prophase I-oocytes were thawed in pre-warmed serial solutions of 1M sucrose in M-199 HEPES Buffered Medium thawing solution (TS) for 60 seconds and then 0.5M sucrose in M-199 HEPES Buffered Medium dilution solution (DS) for 3 minutes. Then, oocytes were recovered in M-199 HEPES Buffered Medium washing solution (WS) for 10 minutes. The oocytes were then moved to pre-equilibrated G-IVF Plus medium (Vitrolife, #10136) supplemented with an additional 10% SPS protein (SAGE, #ART-3011) and incubated at 6.5% CO2 in air at 37°C for 3 hours. The oocytes were then moved to pre-equilibrated GIVF Plus culture drops for continued culture at 6.5% CO2 in air at 37°C.

Oocytes were incubated in pre-equilibrated G-IVF Plus media with the respective inhibitor treatment for 15 hours. Oocytes were fixed in 4% Paraformaldehyde in PHEM buffer 1X supplemented with 0.25% Triton X-100 (Sigma, #1001124827) for 30 minutes at room temperature. The oocytes were transferred to permeabilization solution (PBS containing 0.25% (vol/vol) Triton X-100) for 15 minutes at room temperature. Oocytes were washed three times PBS containing 0.05% Tween-20 and blocked in (0.3% BSA containing 0.05% Tween in PBS) for 1 hour at room temperature. After blocking, the oocytes were incubated in primary antibody in a dark and humidified chamber for overnight at 4C, followed for 3 washes in washing solution of 10 minutes each; then oocytes were incubated in secondary antibody in a dark and humidified chamber for 1 hour at 37C, followed for 3 washes in washing solution of 10 minutes each. Lastly, oocytes were mounted in Vectashield (Vector Laboratories, #H-1000) supplemented with 4′, 6-Diamidino-2-Phenylindole, Dihydrochloride (DAPI; Life Technologies #D1306; 1:170).

### Plasmid information

Preparation of pYX-AURKA-EYFP, pYX-AURKA-EYFP-CDK5RAP2-MBD plasmids were described previously (Blengini et al 2024). The mouse GFP-HEC1 WT plasmid was a gift from Dr. Iain Cheeseman (MIT/Whitehead Institute), mHEC1 cDNA (Imageclone 3709641) was amplified by PCR using the following primers: 5’-GCGCGTCTAGAATGAAGCGCAGTTCAGTTTCCAC-3’ and 5’-TGCTGCCGCGGCATTTGTCGGGAGCCTTAAGTTG-3’ in a backbone with T7 and Xenopus globin UTRs. We mutated the Serine 69 to Alanine (69A) or aspartic acid (69D) using the QuickChange II Site-directed Mutagenesis kit (Agilent Technologies, #200523). The primer for mutating S69-A was: 5’-TAGCGGACATGGATCCAGGAAT**<u>GCT</u>**CAACTTGGTATATTTTCC-3’ and the primer for mutating S69-D was 5’-CTAGCGGACATGGATCCAGGAAT**<u>GAT</u>**CAACTTGGTATATTTTCCA-3’ For *in vitro* transcription, we linearized the plasmids using the Nde1 restriction enzyme and used an mMESSAGE machine T7 kit to generate RNA (Invitrogen, #AM1344). All mRNAs were purified using SeraMag Speedbead (Sigma Aldrich, #GE65152105050250) and stored at - 80°C.

### Immunofluorescence

Oocytes were fixed in 2% Paraformaldehyde in PHEM buffer 1X for 20 minutes at room temperature. Immunofluorescence was performed as previously described (Blengini and Schindler, 2018). Briefly, the oocytes were transferred to permeabilization solution (PBS containing 0.1% (vol/vol) Triton X-100 and 0.3% (wt/vol) BSA) for 20 minutes at room temperature and blocked in (0.3% BSA containing 0.01% Tween in PBS) for 10 minutes. After blocking, the oocytes were incubated in primary antibody in a dark and humidified chamber for 2 hours at room temperature, followed for 3 washes in blocking solution of 10 minutes each; then oocytes were incubated in secondary antibody in a dark and humidified chamber for 1 hour at room temperature, followed for 3 washes in blocking solution of 10 minutes each. Lastly, oocytes were mounted in Vectashield (Vector Laboratories, #H-1000) supplemented with 4′, 6-Diamidino-2-Phenylindole, Dihydrochloride (DAPI; Life Technologies #D1306; 1:170).

### Western Blotting

50 metaphase I oocytes were lysed with Laemmli buffer (Bio-Rad, #161-0737) and denatured at 95C for 10 min. Proteins were separated by electrophoresis in 10% SDS polyacrylamide precast gels (Bio-Rad, #456-1036). The separated polypeptides were transferred to nitrocellulose membrane (Bio-Rad, #170-4156) using a Trans Blot Turbo Transfer System (Bio-Rad) and then blocked with 5% ECL blocking (Amersham, #RPN418) solution in TBS-T (Tris-buffered saline with 0.1% Tween 20) for at least 1h. The membrane was incubated overnight with anti-GFP (Rabbit, 1:500; Sigma, #G1544), or 1 h with anti-Tubulin (Rabbit, 1:1000; Cell Signaling Technology, 11H10) as a loading control. After washing with TBS-T five times, the membranes were incubated with anti-rabbit secondary antibody (1:1000; Kindle Bioscience, #R1006) for 1 h followed with washing with TBS-T five times. The signals were detected using the ECL Select western blotting detection reagents (Kindle Bioscience, #R1002) following the manufacturers protocol. Membranes were stripped prior to loading control detection using Blot Stripping Buffer (ThermoFisher Scientific #46430) for 30 minutes at room temperature.

### Antibodies and drugs

To inhibit the Aurora kinases we used small molecular inhibitors: MLN8237 (AURKA) (Alisertib, Selleckchem #S1133) at 1uM and AZD1152 (AURKB/C) (Selleckchem, #S1147) at 100nM based on previous studies (Blengini et al., 2022). Primary antibodies used were: phosphorylated HEC1 at serine 44 (pSer44) and phosphorylated HEC1 at Serine 69 (pSer69) (DeLuca et al., 2011; DeLuca et al., 2018)(rabbit, 1:750; Gifts from Dr. Jennifer De Luca, Colorado State U.); phosphorylated HEC1 at serine 55 (pHEC1 s55) (rabbit; 1:100; Genetex, #GTX70017); ACA (1:30, Antibodies Incorporated #15-234), TACC3 (Rabbit, 1:100; Novus Biologicals # NBP2-67671), and Alpha-Tubulin (Sheep, 1:100; Cytoskeleton #ATN02). Secondary antibodies were used at 1:200 for IF experiments: anti-mouse-Alexa-568 (Life Technologies #A10037), anti-rabbit-Alexa-647 (Life Technologies #A31573) and Anti-sheep-Alexa-488 (Life Technologies #A11015)

### Microscopy

Images were captured using a Leica SP8 confocal microscope equipped with 40X. 1.30 NA oil immersion objective. Optical Z-stacks of 0.5 µm step with a zoom of 4.5. In those experiments where pixel intensity was compared, the laser power was kept constant among genotypes or treatments.

### Image analysis

For pixel intensity at kinetochores, images were analyzed using ImageJ software (NIH) (Schneider et al., 2012). We created maximum z-projections of the z-sections of each oocyte. To measure kinetochore pixel intensity of phosphorylated HEC1, ACA was used as a mask to define the region of interest. The threshold algorithm was set in WT or control oocytes. At least 20 individual kinetochores per oocyte were measured and the average intensity for each oocyte was calculated for these 20 measurements. Relative pixel intensity was determined by dividing the average intensity by the average intensity of all WT/ control oocytes in the experiment. The volume of spindle was performed using Imaris software (Bitplane). Chromosome misalignment was defined when at least one chromosome was completed separated from the metaphase plate.

### Quantification and Statistical Analysis

All experiments were conducted 2-3 times, any exception would be clarified in the figure legend. Student T-test, ANOVA one-way or two-way analysis were used to evaluate significant differences between/among groups, using Prism software (GraphPad software). Data is shown as the mean ± standard error of the mean (SEM).

## Supplementary information

**Figure S1. Compensatory ability of AURKs to phosphorylate Serines 55 and 69.**

(A) Representative confocal images of oocytes from the indicated genotype at metaphase I and stained with phospho-specific antibodies to detect pSer55 (A) and pSer69 (C) (gray), ACA to detect kinetochores (red) and DAPI to detect DNA (blue). (B), (D), Quantification of phosphorylated HEC1 at pSer55 (B) (purple dots) in (A) and pSer69 (D) (blue dots) in (C). One-way ANOVA: **** p<0.0001; *** p<0.001, ** p>0.01; see complete statistical analysis in Table S1. Scale bar: 10µm.

**Figure S2. Targeting AURKA to MTOCs rescues HEC1 phosphorylation at Ser69 but not at Ser55.**

Representative confocal images of Metaphase I ABC-KO oocytes expressing the indicated constructs. Non-injected ABC-KO oocytes were control. Oocytes were stained with pSer55 (gray) and DAPI (blue). Localization of the targeted AURKA and HEC1 constructs are in gray. (B) Representative confocal images of Metaphase I ABC-KO oocytes expressing the indicated constructs. Non-injected ABC-KO oocytes were control. Oocytes were stained with pSer69 antibodies (gray), ACA (pink) and DAPI (blue). Localization of the targeted AURKA and HEC1 constructs are in gray. (C), (D), (E) Quantification of the relative pixel intensity of pSer69 from (B). Statistical analysis compared to non-injected oocytes is shown (C), to oocytes expressing WT-AURKA is shown in (D) and the rest of the comparisons is shown in (E). One-way ANOVA, **** p<0.0001. Scale bar: 10µm. The number of oocytes analyzed is in brackets.

## Acknowledgments

This work was funded by NIH grant R35 GM136340 to KS. The authors acknowledge Dr. Jennifer DeLuca and Dr. Iain Cheeseman for generously shared reagents and feedback on the project and manuscript. The authors acknowledge members of the Schindler lab for helpful discussions.

## Author contributions

Conceptualization, C.S.B and K.S.; experimental execution: C.S.B; methodology, C.S.B., R.J.M.; providing human oocytes: R.J.M., G.J.G, AND J.E.S; data analysis, C.S.B.; visualization, C.S.B.; writing original draft, C.S.B. and K.S.; review and editing article, C.S.B, R.J.M., G.J.G, AND J.E.S and K.S.; funding, K.S.

