## Supplementary figures and images for "AURKA controls oocyte spindle assembly checkpoint and chromosome alignment by HEC1 phosphorylation"

### Supplemental Figure 1

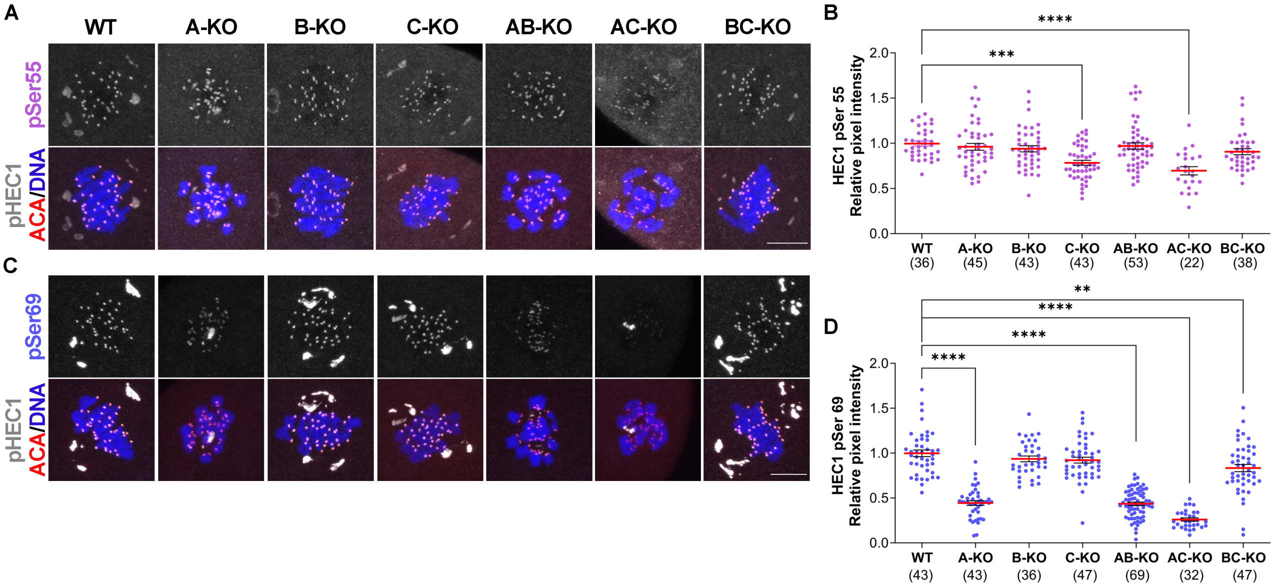

### Supplemental Figure 2

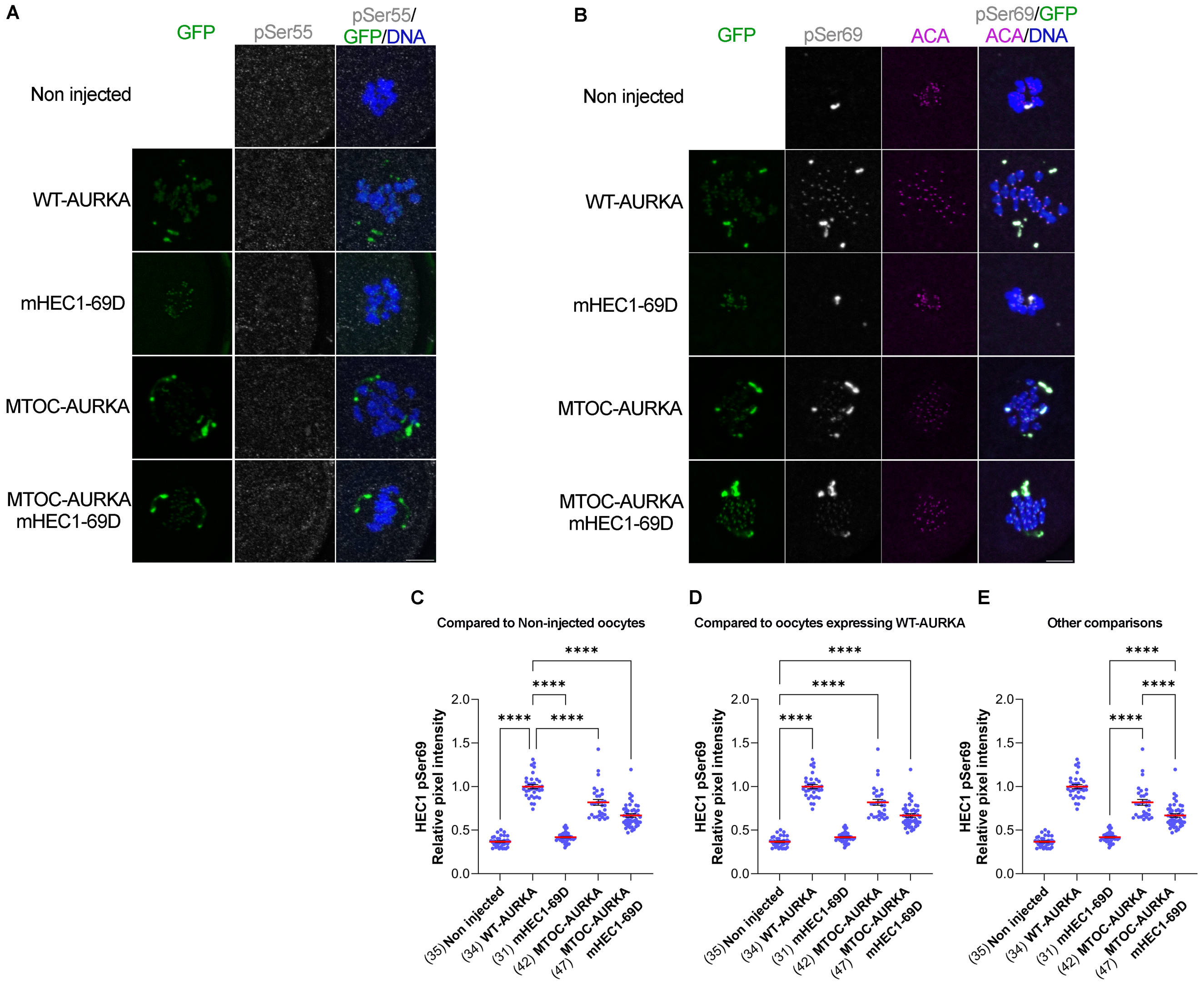
