## Supplemental Table 1 for "AURKA controls oocyte spindle assembly checkpoint and chromosome alignment by HEC1 phosphorylation"

**Table S1**. Complete statistical analysis comparing phosphorylation of different sites of HEC1 between AURK deficient oocytes. One-way ANOVA for each phosphorylation site.

| Comparison | Phosphorylation site | |
|---|---|---|
| | pSer55 | pSer69 |
| WT vs. A-KO | n.s. | **** |
| WT vs. B-KO | n.s. | n.s. |
| WT vs. C-KO | *** | n.s. |
| WT vs. AB-KO | n.s. | **** |
| WT vs. AC-KO | **** | **** |
| WT vs. BC-KO | n.s. | ** |
| A-KO vs. B-KO | n.s. | **** |
| A-KO vs. C-KO | ** | **** |
| A-KO vs. AB-KO | n.s. | n.s. |
| A-KO vs. AC-KO | *** | ** |
| A-KO vs. BC-KO | n.s. | **** |
| B-KO vs. C-KO | * | n.s. |
| B-KO vs. AB-KO | n.s. | **** |
| B-KO vs. AC-KO | *** | **** |
| B-KO vs. BC-KO | n.s. | n.s. |
| C-KO vs. AB-KO | *** | **** |
| C-KO vs. AC-KO | n.s. | **** |
| C-KO vs. BC-KO | n.s. | n.s. |
| AB-KO vs. AC-KO | **** | ** |
| AB-KO vs. BC-KO | n.s. | **** |
| AC-KO vs. BC-KO | ** | **** |
